# Effects of alcohol on gut microbiome in adolescent and adult MMTV-Wnt1 mice

**DOI:** 10.1101/2025.03.17.643801

**Authors:** Hui Li, Leeann Aguilar Meza, Shailesh K Shahi, Zuohui Zhang, Wen Wen, Di Hu, Hong Lin, Ashutosh Mangalam, Jia Luo

**Affiliations:** Department of Pathology, University of Iowa Carver College of Medicine, Iowa City, IA 52242, USA; Holden Comprehensive Cancer Center, University of Iowa, Iowa City, IA, 52242, USA

**Author notes:** Correspondence author: Jia Luo, Department of Pathology, University of Iowa Carver College of Medicine, Iowa City, IA 52242, USA.

**Keywords:** Alcohol misuse, breast cancer, gut dysbiosis, tumor promotion

## Abstract

Breast cancer is the most commonly diagnosed cancer in women worldwide, with alcohol consumption recognized as a significant risk factor. While epidemiological studies consistently show a positive correlation between alcohol consumption and increased breast cancer risk, the underlying mechanisms remain unclear. Recent evidence suggests that the gut microbiome—the diverse collection of microorganisms, including bacteria, viruses, and fungi, residing in the gastrointestinal tract—plays a pivotal role in systemic health and disease. This is achieved through its regulation of key physiological processes such as metabolism, immune function, and inflammatory responses. Disruption of the gut microbiome (dysbiosis) has recently been implicated in the development of breast cancer. We hypothesized that alcohol exposure induces gut dysbiosis, which in turn drives systemic inflammation and carcinogenic processes. Previously, we demonstrated that alcohol exposure promotes mammary tumor growth and aggressiveness in MMTV-Wnt1 (Wnt1) transgenic mice, an established model for investigating mechanisms of alcohol-induced tumor promotion. In this study, we sought to determine whether alcohol exposure induces gut dysbiosis in adolescent and adult Wnt1 transgenic mice and their wild-type FVB counterparts. Our findings revealed that alcohol exposure significantly reduced microbiome richness in adult Wnt1 and FVB mice. Alcohol exposure also markedly altered microbiome composition in adolescents and adults in both strains. Additionally, we identified specific microbial taxa that were significantly affected by alcohol exposure. These results demonstrate that alcohol disrupts the gut microbiome in a preclinical breast cancer model, providing insights into the potential role of gut dysbiosis in alcohol-induced mammary tumor promotion and offering avenues for future research.

## Introduction

Alcohol consumption has emerged as a critical risk factor for breast cancer as many epidemiological and experimental studies have demonstrated a positive correlation between alcohol consumption and increased breast cancer risk (Chen, Rosner et al. 2011, Seitz, Pelucchi et al. 2012, McDonald, Goyal et al. 2013, Bagnardi, Rota et al. 2015, Wang, Xu et al. 2017, LoConte, Brewster et al. 2018). However, the cellular and molecular mechanisms underlying alcohol’s tumor promotion remain unclear. There are several proposed mechanisms. For example, alcohol consumption can elevate the estrogen levels in the both premenopausal and postmenopausal women, which may contribute to the alcoholic effects on the higher breast cancer risk (Reichman, Judd et al. 1993, Gavaler, Galvao-Teles et al. 2002, Register, Cline et al. 2002, Tin Tin, Smith-Byrne et al. 2024). Alcohol and its metabolite, acetaldehyde, are both known to damage DNA and induce gene mutation (Dumitrescu and Shields 2005, Liu, Nguyen et al. 2015). Alcohol exposure can also promote the accumulation of excessive reactive oxygen species (ROS) and oxidative stress, which may promote mammary carcinogenesis and aggressiveness (Dumitrescu and Shields 2005, Wang, Xu et al. 2017). Recently, studies have shown that alcohol consumption may also affect gut microbiome, an essential regulator of systemic inflammation, estrogen metabolism, and immune responses, suggesting a *novel* pathway through which alcohol may impact breast cancer risk and progression (Bajaj 2019, Addolorato, Ponziani et al. 2020, Koutromanos, Legaki et al. 2024).

Gut microbiome is the complex community of microbes such as bacterial, viruses and fungi that reside in the gastrointestinal system and modulate the functions of local and distant organs through metabolic, immunologic and hormonal pathways (Brestoff and Artis 2013, Sommer and Bäckhed 2013, Ruff and Kriegel 2015). A disruption in the composition of gut microbiome, known as gut dysbiosis, is characterized by the reduced microbial diversity, loss of beneficial bacteria or the overgrowth of harmful bacteria. Gut dysbiosis has been linked to a wide range of diseases including breast cancer (Yang, Tan et al. 2017, Zhang, Xia et al. 2021). Recent studies have suggested that gut dysbiosis play a role in various aspects of breast cancer, including tumorigenesis, disease progression, metastasis and treatment outcome (Laborda-Illanes, Sanchez-Alcoholado et al. 2020, Ruo, Alkayyali et al. 2021, Terrisse, Derosa et al. 2021, Zhang, Xie et al. 2022). For example, a gut microbiome profiling study conducted the Midwest United States revealed gut dysbiosis in breast cancer patients, characterized with the depletion of short-chain fatty acid-producing gut bacteria (Shrode, Knobbe et al. 2023). Additionally, a pilot study reported associations between gut microbiome composition and breast tumor characteristics, such as receptor status, stage, and grade, as well as established breast cancer risk factors (Wu, Tseng et al. 2020).

Emerging evidence has demonstrated that alcohol consumption can disrupt gut microbiome and alcohol-induced gut dysbiosis is considered an early factor in alcohol-related disorders such as alcohol use disorders (AUD) and alcoholic-liver disease (ALD) (Mutlu, Keshavarzian et al. 2009, Bajaj 2019, Addolorato, Ponziani et al. 2020, Koutromanos, Legaki et al. 2024). The role of gut dysbiosis in alcohol-related breast cancer, however, has not yet been studied. Using a mammary tumorigenesis model of MMTV-Wnt1 transgenic mice, we have previously showed that alcohol exposure enhanced tumorigenesis and aggressiveness, with adolescent mice showing greater sensitivity to alcoholic effects than adults (Xu, Li et al. 2023). In this study, we aimed to determine whether alcohol exposure alters gut microbiome in MMTV-Wnt1 mice prior to the onset of mammary tumorigenesis by analyzing changes in gut bacterial composition following alcohol exposure.

## Material and methods

### Animals and experimental groups

FVB MMTV-Wnt1 [FVB.Cg-Tg (Wnt1)1Hev/J] transgenic and FVB *wild type* (*WT*) mice were obtained from The Jackson Laboratories (Bar Harbor, ME), bred, and housed in a climate-controlled animal facility. All procedures were reviewed and approved by the Institutional Animal Care and Use Committee (IACUC) of the University of Iowa. In this study only female mice were used. Adolescent mice (5-week-old) or adult mice (10-week-old) from either FVB *wt* or MMTV-Wnt1 (Wnt1) transgenic strain were assigned into control and alcohol exposure groups. Two ages were selected because we previously demonstrated that adolescent Wnt1 mice were more susceptible to alcohol-induced mammary tumor promotion than adult mice (Xu et al., 2023). For alcohol exposure, the animals received a daily intraperitoneal (IP) injection of either PBS (control) or ethanol solution (2.5 g/kg, 25% w/v) for 15 days. The acute exposure regime was chosen to model binge alcohol exposure (Xu, Li et al. 2023). All mice were monitored daily by palpation to ensure none developed mammary tumor.

The experimental groups were assigned based on the age, strain, and treatment (Figure 1). The adolescent mice started at 5 weeks old (5W) and became 7 weeks old (7W) after 15 days of treatment. The adult mice started at 10 weeks old (10W) and became 12 weeks old (12W) after 15-day treatment period. There were four experimental groups of animals used in this study: 1) Adolescent FVB which includes control group before (Control_5W, n = 7) or after PBS treatment (Control_7W, n = 7) and ethanol group before (Ethanol_5W, n = 5) or after ethanol treatment (Ethanol_7W, n = 5); 2) Adult FVB including control group before (Control_10W, n = 8) or after PBS treatment (Control_12W, n = 8) and ethanol group before (Ethanol_10W, n = 8) or after ethanol treatment (Ethanol_12W, n = 8); 3) Adolescent Wnt1 which includes control group before (Control_5W, n = 5) or after PBS treatment (Control_7W, n = 5) and ethanol group before (Ethanol_5W, n = 7) or after ethanol treatment (Ethanol_7W, n = 7); 4) Adult Wnt1 including control group before (Control_10W, n = 6) or after PBS treatment (Control_12W, n = 6) and ethanol group before (Ethanol_10W, n = 7) or after ethanol treatment (Ethanol_12W, n = 7).

**Figure 1:**
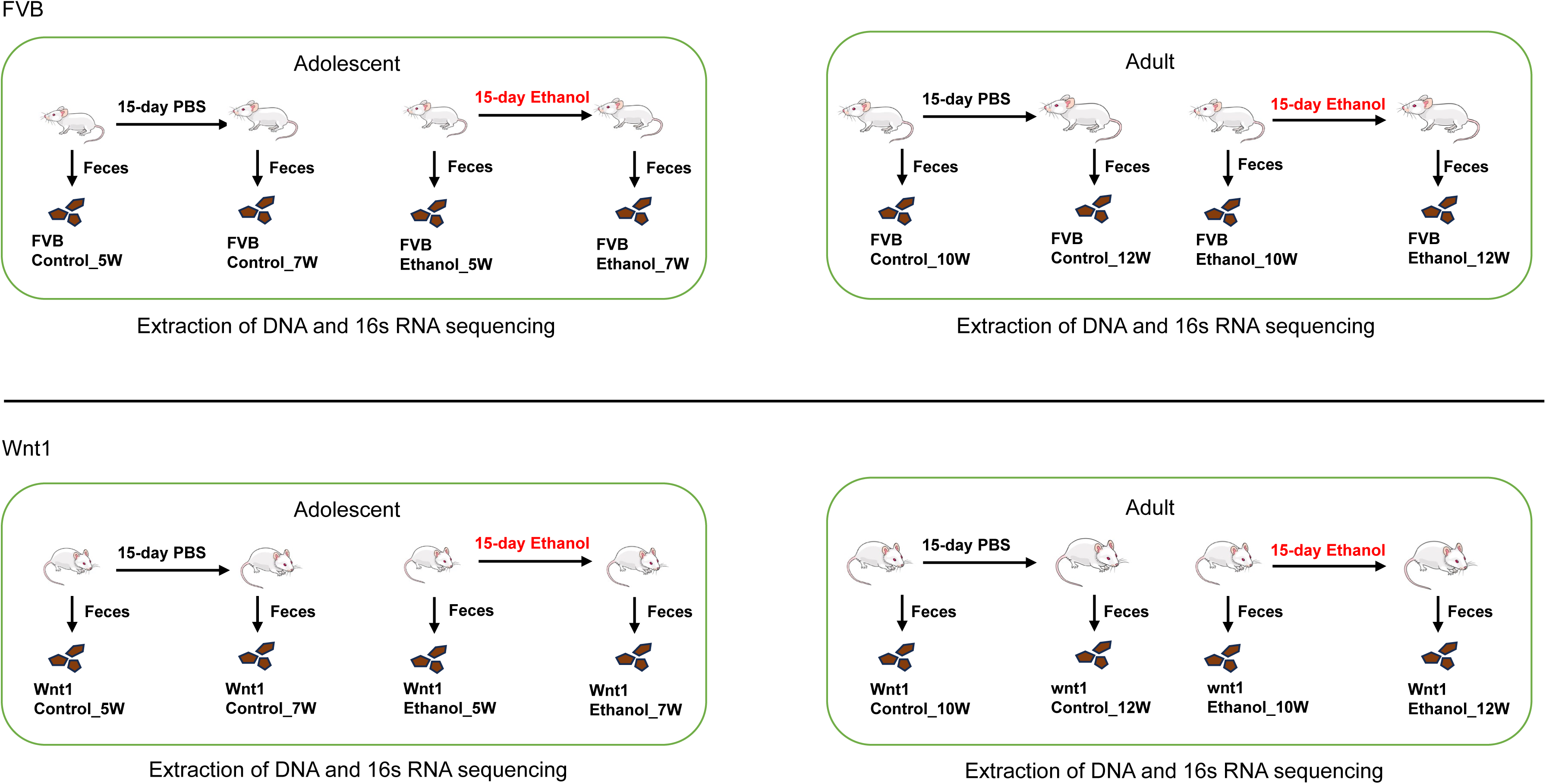
Design of experimental groups. Both adolescent (5-week-old) and adult (10-week-old) FVB and Wnt1 mice were treated with either PBS or ethanol for 15 days and one day after the treatment, their feces were collected and sent for DNA purification and 16s rRNA sequencing for gut microbiome composition. The number of each experimental groups are as follows: Adolescent FVB which includes control group before (FVB control_5W, n = 7) or after PBS treatment (FVB control_7W, n = 7) and ethanol group before (FVB Ethanol_5W, n = 5) or after ethanol treatment (FVB Ethanol_7W, n = 5); 2) Adult FVB including control group before (FVB control_10W, n = 8) or after PBS treatment (FVB control_12W, n = 8) and ethanol group before (FVB Ethanol_10W, n = 8) or after ethanol treatment (FVB Ethanol_12W, n = 8); 3) Adolescent Wnt1 which includes control group before (Wnt1 control_5W, n = 5) or after PBS treatment (Wnt1 control_7W, n = 5) and ethanol group before (Wnt1 Ethanol_5W, n = 7) or after ethanol treatment (Wnt1 Ethanol_7W, n = 7); 4) Adult Wnt1 including control group before (Wnt1 control_10W, n = 6) or after PBS treatment (Wnt1 control_12W, n = 6) and ethanol group before (Wnt1 Ethanol_10W, n = 7) or after ethanol treatment (Wnt1 Ethanol_12W, n = 7).

### Fecal sample collection, extraction of DNA and 16s RNA sequencing

Fecal samples from each animal were collected either one day before or after the exposure and stored at -80°C freezer until further processing for DNA extraction. Microbial DNA extraction, 16S rRNA amplicon, and sequencing were performed as previously published protocol (Shahi, Zarei et al. 2019). Briefly, DNA was isolated using DNesay PowerLyzer PowerSoil Kit (Qiagen, Germantown, MD) as per the manufacturer’s instructions, including the recommended bead-beating step. The sequencing library was prepared using a 2-step amplification, where the V3-V4 region of bacterial 16s rRNA gene was amplified in step 1, and barcodes were added using Nextera XT Index Kit (Illumina MiSeq) in step 2. PCR products were purified and sequenced using Illumina MiSeq platform. R based platform Divisive Amplicon Denoising Algorithm 2 (DADA2) (Callahan, McMurdie et al. 2016)was used to trim, merge, and filter reads and generate an amplicon sequence variant (ASV) table. The ASVs were taxonomically classified from kingdom to species levels using the Silva database (version 138.1), with a median read count of 49,839 (ranging from 1,667 to 78,455 reads).

### Microbiome analyses and visualization

Microbial communities were analyzed using previously described methods for each experimental group (Shahi, Zarei et al. 2019, Yadav, Ali et al. 2022, Shrode, Knobbe et al. 2023). Briefly, custom R (Version 4.3.1) scripts were utilized, integrating packages such as phyloseq (McMurdie and Holmes 2013), vegan (Oksanen J 2024), ggpubr (A 2023), dplyr (Wickham H 2023), microbiome (Lahti L 2017), tidyr (Wickham H 2024), sigminer (Wang S 2024), and ggplot2 (Wickham H 2016). Except for alpha diversity, reads underwent normalization using constant-sum scaling and log10 transformation at the bacterial level to their median sequencing depth. Alpha diversity analysis was conducted on unfiltered data using the Chao1 index. Beta diversity was assessed via Principal Component Analysis (PCA) based on weighted UniFrac distances, with significance tested through PERMANOVA. A heatmap of the top most abundant genera was generated using the phyloseq (McMurdie and Holmes 2013) and ggplot2 (Wickham H 2016) packages, visualizing top bacterial genera based on weighted UniFrac distances (Lozupone and Knight 2005). Multidimensional Scaling (MDS) was employed for ordination, with sample groups arranged along the x-axis to represent relative abundance. To visualize enrichment, the LEfSe plot was produced using the microbiomeMarker (Cao, Dong et al. 2022) package’s “run_lefse” function, highlighting the genera enriched in the different groups within each experimental group using the Kruskal-Wallis test.

### Statistical analyses

A two-way analysis of variance (ANOVA) was conducted with age and treatment as independent variables to analyze the relative abundance of selected microbial taxa across experimental groups. Post hoc comparisons were performed using Tukey’s correction for multiple comparisons, utilizing GraphPad Prism Version 10.3.1 (GraphPad Software, Inc., www.graphpad.com). Additionally, the Wilcoxon matched-pairs signed rank test was applied to assess the effects of alcohol on microbial taxa abundance in selective treatment groups. A significance threshold of p < 0.05 was set for all analyses.

## Results

### Microbiome analysis of alpha diversity of alcohol-exposed mice

Alpha diversity is within sample diversity. When examining alpha diversity, we are able to evaluate the distribution of microbes within a sample or metadata category. Chao1 index, a statistical estimator that measures species richness, is widely used to assess alpha diversity in microbiome research including gut microbiome (Jensen, Cady et al. 2021). We used Chao1 index to determine the effect of alcohol exposure on alpha diversity (Figure 2). In FVB mice, there was no significant difference in the Chao1 index between control groups in either adolescents (p _control_7W vs. control_5W_ = 0.16) or adults (p_control_12W vs. control_10W_ = 0.14). Alcohol exposure did not significantly change the Chao1 index in adolescent mice (p_Ethanol_7W vs. Ethanol_5W_ = 0.2), but significantly reduced the number of microbial species in adults (p_Ethanol_12W vs. Ethanol_10W_ = 0.0013). In Wnt1 mice, the Chao1 index similarly showed no significant differences between control groups either in adolescents (p_control_7W vs. control_5W_ = 1) or adults (p_control_12W vs. control_10W_ = 1). Alcohol exposure did not significantly alter the Chao1 index in adolescents (p_Ethanol_7W vs. Ethanol_5W_ = 0.28), but significantly reduced microbial richness in adults (p_Ethanol_12W vs. Ethanol_10W_ = 0.0023). These findings demonstrated that alcohol exposure significantly reduced alpha diversity in adults in both mouse strains, whereas the impact on the adolescents was not significant.

**Figure 2:**
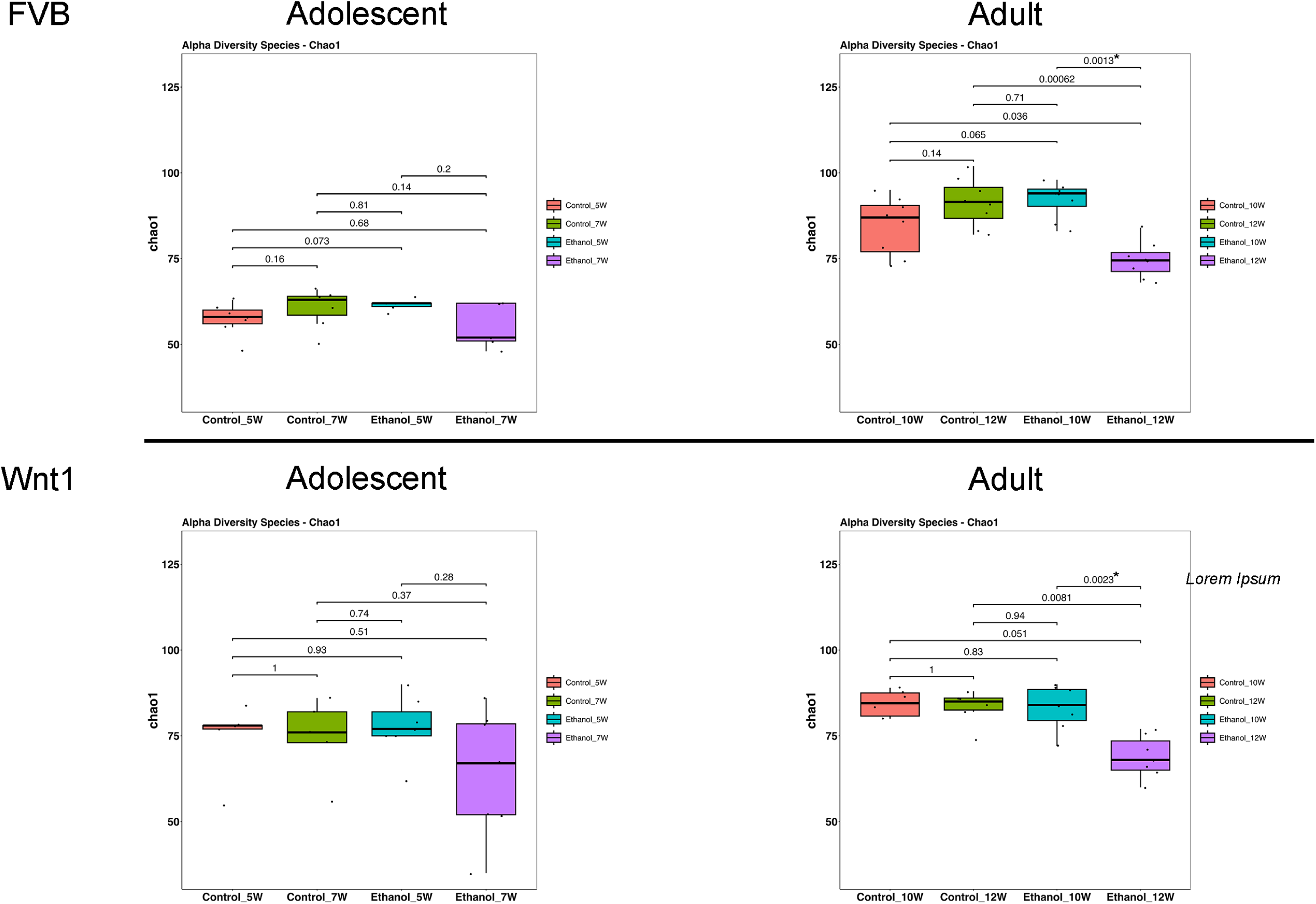
Effects of alcohol on microbial species richness. The effect of alcohol on alpha diversity was measured by Chao1 index in all experimental groups. Alcohol exposure significantly reduced alpha diversity in adult mice.

### Microbiome analysis of beta diversity of alcohol-exposed mice

Beta diversity is the diversity between samples and a common statistical method to assess the similarity or differences in microbial compositions between samples (Jensen, Cady et al. 2021). To examine the impact of alcohol exposure on microbial diversity across experimental groups, we employed weighted UniFrac, a quantitative measure of beta diversity (Figure 3). In FVB mice, treatment significantly affected gut microbial composition in both adolescents (p = 0.001) and adults (p = 0.001). Further analysis by treatment revealed significant differences in the microbial community in controls across age groups for both adolescents (p_control_7W vs. control_5W_ = 0.01) and adults (p_control_12W vs. control_10W_ = 0.025), and an even more marked differences in alcohol groups in both adolescents (p_Ethanol_7W vs. Ethanol_5W_ = 0.002) and adults (p_Ethanol_12W vs. Ethanol_10W_ = 0.002). A similar trend was observed in Wnt1 mice, with a near-significant treatment effect on gut microbial composition in adolescents (p = 0.076), and a significant effect in adults (p = 0.013). When separated by treatment, the control groups showed a nearly significant difference in adolescents (p_control_7W vs. control_5W_ = 0.055) and adults (p_control_12W vs. control_10W_ = 0.088), while significant differences were observed in alcohol-exposed adolescents (p_Ethanol_7W vs. Ethanol_5W_ = 0.013) and adults (p_Ethanol_12W vs. Ethanol_10W_ = 0.014). These results suggested that alcohol exposure has a more pronounced impact on the beta diversity, compared to the effects with natural age-related development in controls regardless of stains and ages.

**Figure 3:**
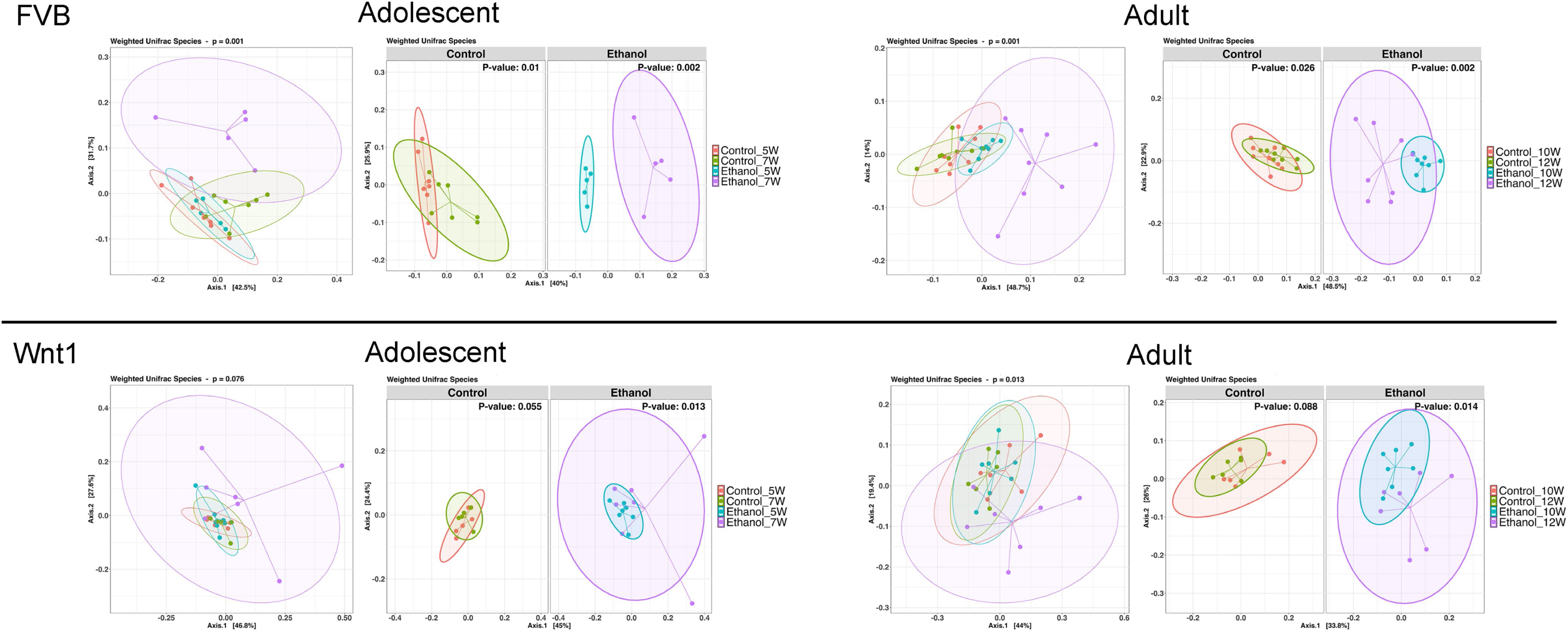
Effects of alcohol on microbial compositions. The effect of alcohol on beta diversity was determined by weighted UniFrac analysis in all experimental groups. Alcohol exposure significantly changed the microbial compositions in adolescents and adults in both strains.

### The abundance of microbial populations of alcohol-exposed mice

To highlight the most abundant microbial taxa in each experimental group, we used heatmap visualization to display the top 20 most prevalent microbial populations (Figure 4). The heatmap analysis revealed a similar pattern of microbial abundance across the experimental groups. In both adolescent and adult FVB experimental groups, the commonly identified taxa included the species *Akkermansia muciniphila*; the genera *Turicibacter*, *Lachnoclostridium*, *Lactobacillus*, *Lachnospiraceae_NK4A136_group*, and *Alistipes*; the families *Muribaculaceae*, *Oscillospiraceae*, and *Ruminococcaceae*; and the order *Clostridia_vadinBB60_group*. In the adolescent and adult Wnt1 experimental groups, the taxa commonly identified were the species *A. muciniphila* and *L. intestinalis*; the genera *Bacteroides*, *Lachnospiraceae_NK4A136_group*, *Lactobacillus*, *Prevotellaceae_UCG-001*, *Alistipes*, *Rikenellaceae_RC9_gut_group*, and *Ruminococcus*; the families *Lachnospiraceae* and *Ruminococcaceae*. The analysis showed distinct yet overlapping microbial profiles across experimental groups, with certain taxa appearing as key microbial populations such as *A. muciniphila*, and members of *Lachnospiraceae* and *Ruminococcaceae* families, present in both FVB and Wnt1 groups. This combination of both shared and distinct microbial populations between the FVB and Wnt1 groups may relate to their age and genetic backgrounds.

**Figure 4:**
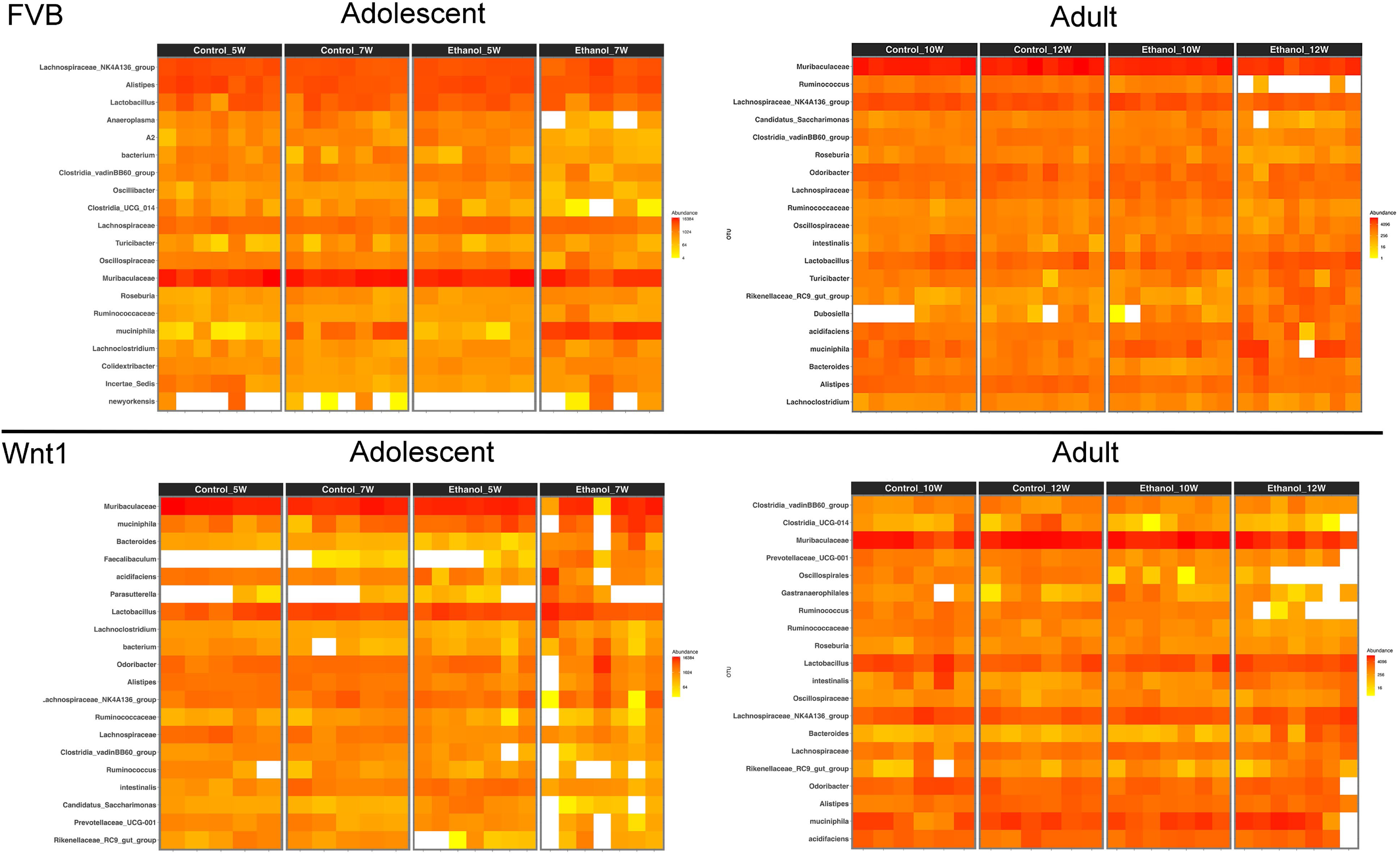
Effects of alcohol on microbial taxa. Top 20 most prevalent microbial taxa in each experimental group were visualized using heat maps.

### The differential microbial populations of alcohol-exposed mice

We then preformed Linear Discriminant Analysis Effect Size (LEfSe) analysis to identify the microbial populations that are mostly affected by alcohol exposure in each experimental group (Figure 5). LEfSe is a biomarker discovery tool that identifies statistically significant differences between multiple groups. In the adolescent FVB experimental group, LEfSe analysis revealed that the species *A. muciniphila* and the genus *Colidextribacter* were the most enriched taxa in alcohol-exposed mice. In the adult FVB experimental group, alcohol exposure was associated with an enrichment of the species *A. muciniphila*, *L. reuteri*, and *P. goldsteinii*, as well as the genera *Bacteroides*, *Dubosiella*, *Rikenellaceae_RC9_gut_group*, and *Parasutterella*. In the adolescent Wnt1 experimental group, alcohol exposure led to a higher abundance of the genera *Faecalibaculum*, *Bacteroides*, and *Turicibacter*. In the adult Wnt1 experimental group, alcohol exposure was associated with an enrichment of the species *P. goldsteinii* and *L. reuteri*, along with the genera *Bacteroides* and *Rikenella*. These results indicated that age and genetic background significantly affect the microbiome’s response to alcohol, with both overlapping and unique taxa affected across experimental groups.

**Figure 5:**
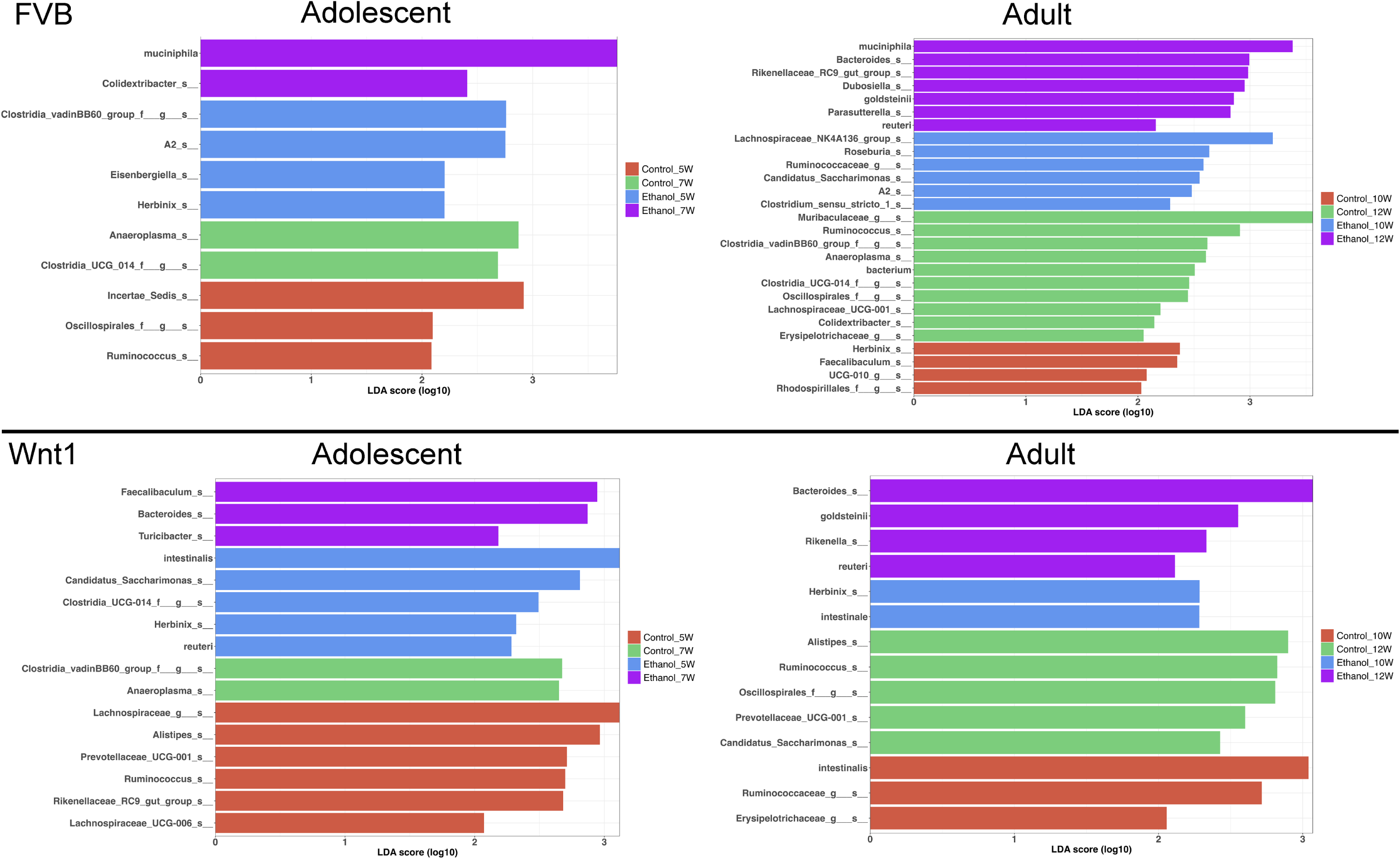
Effects of alcohol on specific microbial taxa. Linear Discriminant Analysis Effect Size (LEfSe) analysis was performed to identify the microbial taxa that were mostly affected by alcohol exposure.

### Comparisons of taxonomic abundance of selective microbial population

We conducted a two-way analysis of variance (ANOVA) for each microbial taxon that was identified in the LEfSe analysis to further validate the findings (data not shown). We further analyzed microbial taxa that were significantly altered by alcohol exposure in adolescent Wnt1 mice, because adolescent Wnt1 mice were more sensitive to alcohol’s tumor promotion compared to adults (Xu, Li et al. 2023). We also compared the relative abundance of specific taxa across other experimental groups.

Alcohol exposure increased the abundance of several microbial taxa in the adolescent Wnt1 experimental group, including *P. goldsteinii* and the genera *Bacteroides, Faecalibaculum,* and *Turicibacter*. The abundance of these taxa was analyzed across various experimental groups using two-way ANOVA analysis followed by Tukey’s post hoc test (Figure 6). *P. goldsteinii* showed a nearly significant increase in alcohol-exposed adolescent Wnt1 mice (p = 0.062) and was significantly enriched in adult Wnt1 and adult FVB mice. Although *Bacteroides* showed a non-significant increase in the adolescent Wnt1 experimental group by two-way ANOVA followed by Tukey’s post hoc test (p = 0.1590), the Wilcoxon test revealed a significant effect (p = 0.0469, Supplementary Figure 1). Both adult Wnt1 and FVB experimental groups exhibited a significant increase in *Bacteroides* following alcohol exposure. *Faecalibaculum* was significantly elevated in alcohol-exposed adolescent Wnt1 mice but was notably decreased in the control group of adult FVB mice. *Turicibacter* showed a significant increase in the adolescent Wnt1 experimental group and a nearly significant increase in the adult FVB experimental group (p = 0.0801).

**Figure 6:**
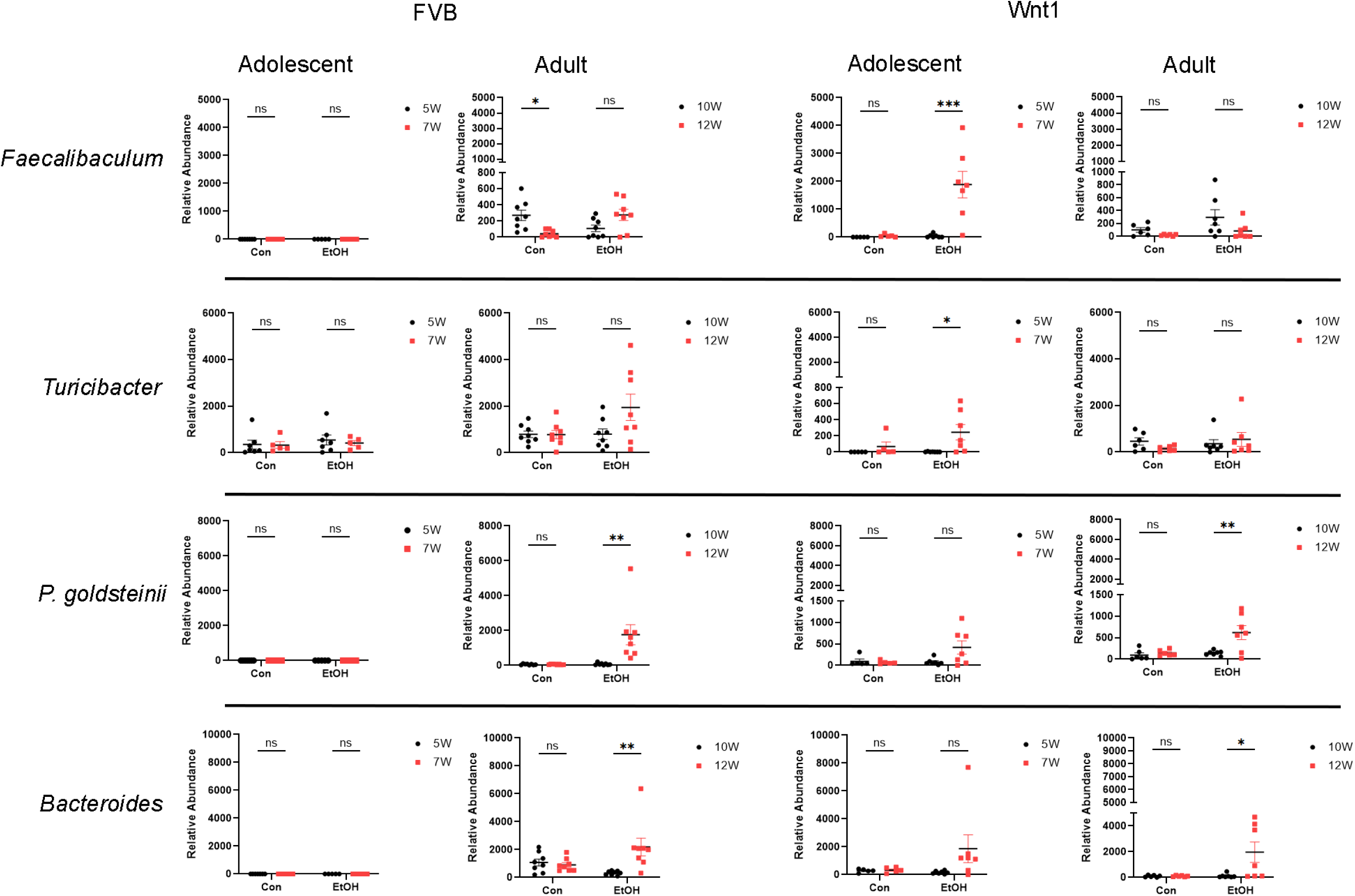
Analysis of alcohol-enriched microbial taxa. Alcohol-enriched microbial taxa were analyzed by two-way ANOVA followed by Tukey’s post hoc test. Selective microbial taxa including the species *P. goldsteinii*; the genera *Faecalibaculum, Turicibacter* and *Bacteroides* were increased by alcohol exposure. FVB: n = 7 for control (Con) 5W and 7W; n = 5 for ethanol (EtOH) 5W and 7W; n = 8 for control (Con) 10W and 12W; n = 8 for ethanol (EtOH) 10W and 12W; Wnt1: n = 5 for control (Con) 5W and 7W; n = 7 for ethanol (EtOH) 5W and 7W; n = 6 for control (Con) 10W and 12W; n = 7 for ethanol (EtOH) 10W and 12W. ∗p < 0.05, ∗∗p < 0.01, ∗∗∗p < 0.001, ns = not significant (p > 0.05).

Although *A. muciniphila* did not show a significant increase in alcohol-exposed adolescent Wnt1 mice, it is one of the most abundant bacterial species in the gut microbiome and was identified in both the heatmap and LEfSe analysis across multiple experimental groups (Supplementary Figure 2). A detailed examination revealed that *A. muciniphila* was significantly elevated by alcohol exposure only in the adolescent FVB group, with no significant changes observed in other groups.

In contrast, the abundance of certain microbial taxa, including the species *L. intestinalis* and the genera *Anaeroplasma*, *Herbinix*, and *Candidatus_Saccharimonas*, was decreased following alcohol exposure in the adolescent Wnt1 experimental group, with further analysis across all experimental groups (Figure 7). *L. intestinalis* was significantly reduced in the alcohol-exposed adolescent Wnt1 experimental group, with no similar effects observed in other experimental groups. In the same experimental group, the *Anaeroplasma* level was significantly higher in control mice but lower in the alcohol-exposed mice, although this reduction was not statistically significant (p = 0.4502). The Wilcoxon test, however, showed a significant effect of alcohol on *Anaeroplasma* (p = 0.0312, Supplementary Figure 1). In addition, *Anaeroplasma* was significantly reduced following alcohol exposure in the adolescent FVB experimental group. *Herbinix* exhibited a consistent decrease following alcohol exposure across the adolescent Wnt1, adult Wnt1, and adolescent FVB experimental groups, with a nearly significant reduction in the adult FVB experimental group (p = 0.0715). Finally, *Candidatus_Saccharimonas* level was decreased in response to alcohol exposure in both the adolescent Wnt1 and adult FVB experimental groups.

**Figure 7:**
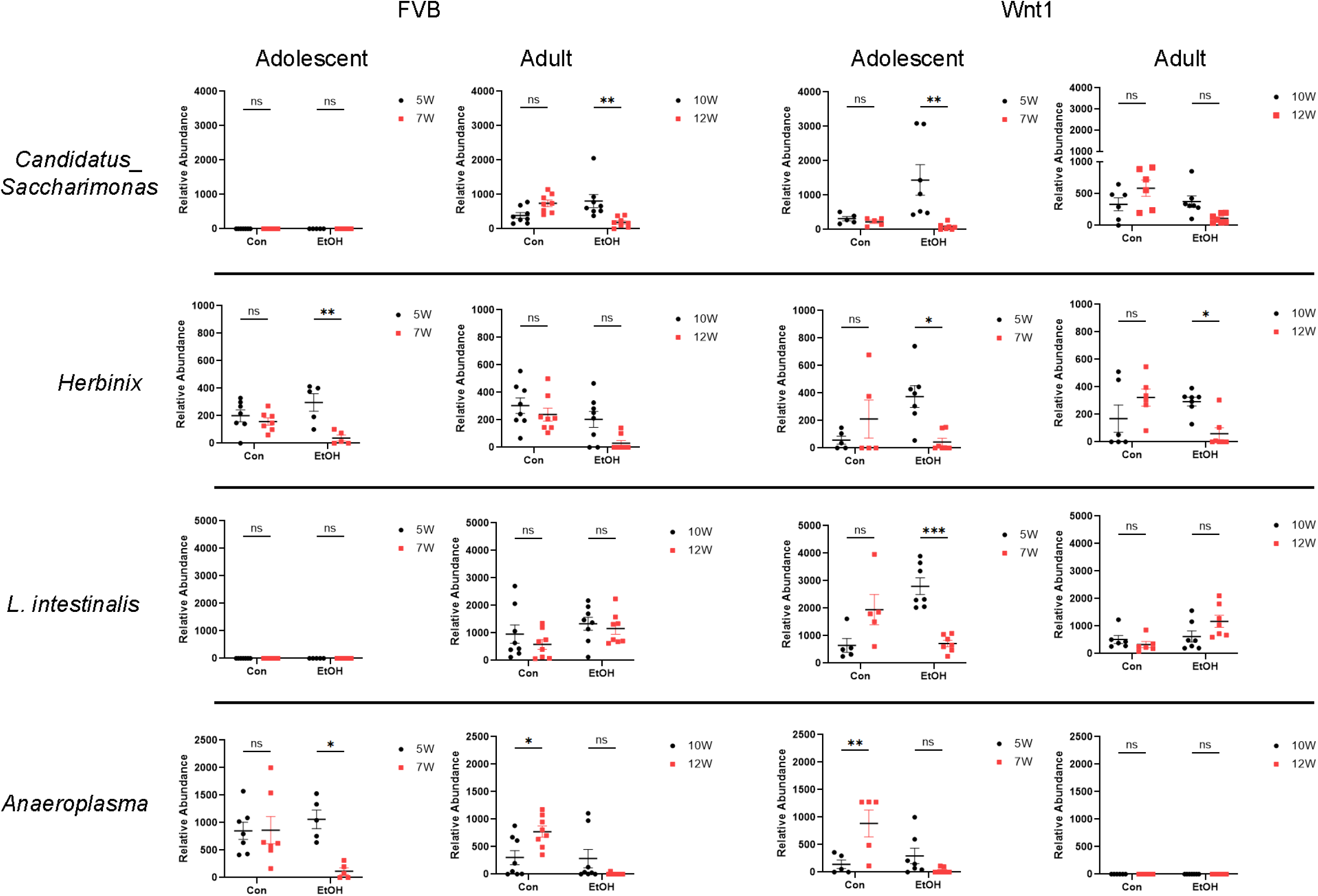
Analysis of alcohol-depleted microbial taxa. Alcohol-depleted microbial taxa were analyzed by two-way ANOVA followed by Tukey’s post hoc test. Selective microbial taxa such as the species *L. intestinalis* and the genera *Candidatus_Saccharimonas, Herbinix* and *Anaeroplasma* were reduced by alcohol exposure. FVB: n = 7 for control (Con) 5W and 7W; n = 5 for ethanol (EtOH) 5W and 7W; n = 8 for control (Con) 10W and 12W; n = 8 for ethanol (EtOH) 10W and 12W; Wnt1: n = 5 for control (Con) 5W and 7W; n = 7 for ethanol (EtOH) 5W and 7W; n = 6 for control (Con) 10W and 12W; n = 7 for ethanol (EtOH) 10W and 12W. ∗p < 0.05, ∗∗p < 0.01, ∗∗∗p < 0.001, ns = not significant (p > 0.05)

## Discussion

The Wnt1 transgene, driven by the mouse mammary tumor virus (MMTV) promoter, is a well-established mouse model to study mammary tumor development, as the Wnt1 signaling pathway is crucial for regulating cell proliferation, differentiation and development (Tsukamoto, Grosschedl et al. 1988, Li, Hively et al. 2000). Previously, we utilized these MMTV-Wnt1 transgenic mice to study alcohol-induced tumor promotion and demonstrated that alcohol exposure shortened the tumor latency, increased lung metastasis, elevated blood levels of estradiol and progesterone, with adolescent mice showing greater sensitivity to the effects of alcohol than adults (Xu, Li et al. 2023). In this study, we further used this model to determine whether alcohol exposure alters the gut microbiome. Our results demonstrated that alcohol exposure significantly reduced microbial species richness as indicated by decreased Chao1 index in adult mice, while it had little effects on this index in adolescent mice. Alcohol also altered microbial compositions as indicated by the analysis of Beta diversity in both adolescents and adults in both strains. Comparative profiling identified a number of taxa consistently affected by alcohol across groups. Further LEfSe and two-way ANOVA analyses confirmed that specific taxa were targets of alcohol exposure.

A lower alpha diversity in the gut microbiome, as indicated by a reduced Chao1 index, has been reported in patients with breast cancer when compared to healthy controls in multiple studies (Goedert, Jones et al. 2015, Ma, Qu et al. 2022, Wu, Vigen et al. 2022, Luan, Ge et al. 2024, Valé, Franck et al. 2024). However, there are studies showing that breast cancer patients have different microbial composition without any difference in alpha diversity (Aarnoutse, Hillege et al. 2021, Yaghjyan, Mai et al. 2021, Shrode, Knobbe et al. 2023). An intriguing finding from our study is that the Chao1 index in adolescent mice appeared more resilient to the adverse effects of alcohol exposure compared to adults (Figure 1), thus suggesting adult gut microbiome may have reduced plasticity in response to environmental factors such as alcohol. The microbiome plasticity, or the ability of the gut microbiome to adapt to environmental changes, is known to be highest early in life and decline with age (Grembi, Nguyen et al. 2020, Qin, Song et al. 2020, Thriene and Michels 2023). This difference in the species richness in response to alcohol exposure between adolescent and adult mice suggests that, while adolescent mice may be more sensitive to alcohol’s action in the context of breast cancer, the effects of alcohol on their gut composition are more complex and multifaceted.

Our beta diversity analysis using weighted UniFrac showed that changes in microbial communities over time were more pronounced in Wnt1 mice compared to FVB mice, with significant alterations observed in the alcohol-exposed groups relative to controls. These results suggest that alcohol exposure significantly disrupts the beta diversity of the gut microbiome over time, beyond the natural developmental changes observed in controls, and Wnt1 mice were more sensitive to alcohol-induced alterations. The increased sensitivity of gut microbiota in Wnt1 mice, may be attributed to the role of Wnt1 signaling in regulating cell proliferation and differentiation, which could interact with gut microbiome, rendering it more susceptible to environmental factors such as alcohol. Notably, altered beta diversity in the gut microbiome has been linked to various stages of breast cancer, including its subtypes and the presence of metastatic disease (Goedert, Jones et al. 2015, Byrd, Vogtmann et al. 2021, Terrisse, Derosa et al. 2021, Wenhui, Zhongyu et al. 2022). These variations in microbial diversity might correlate with systemic inflammation or immune markers, potentially influencing tumor progression and responses to therapies such as chemotherapy or hormone therapy (Jørgensen, Trøseid et al. 2016, Cano-Ortiz, Laborda-Illanes et al. 2020).

Comparative analysis of microbial profiles, visualized as heatmaps, revealed distinct yet overlapping responses to alcohol exposure across experimental groups in both FVB and Wnt1 mice. The commonly identified taxa in all experimental groups included the species *A. muciniphila*; the genera *Lachnospiraceae_NK4A136_group*, *Lactobacillus* and *Alistipes*; and the family *Ruminococcaceae. A. muciniphila* is a mucus-degrading bacterium that resides in the gut’s mucus layer and is often associated with gut barrier integrity and metabolic health (Macchione, Lopetuso et al. 2019, Mo, Lou et al. 2024). The genus *Lachnospiraceae_NK4A136_group*, part of the family of *Lachnospiraceae*, is known for producing short-chain fatty acids (SCFAs) such as butyrate, which support gut health (Hu, Wang et al. 2019, Ma, Ni et al. 2020). The genus *Lactobacillus* comprises numerous probiotic species known for their ability to produce lactic acid and promote gut health (Slover and Danziger 2008). Additionally, *Alistipes*, from the *Bacteroidetes* phylum, is often linked with protein fermentation and bile acid metabolism (Parker, Wearsch et al. 2020). The *Ruminococcaceae* family plays a role in fiber degradation and SCFA production, including butyrate, which supports gut barrier function (Biddle, Stewart et al. 2013). Although Wnt1 signaling can affect the gut microbiome through its effects on the gut environment and epithelial cell turnover (Moparthi and Koch 2019, Liang, Liu et al. 2022), the similar microbial profiles observed in both FVB and Wnt1 mice suggest that alcohol exposure exert a universal impact on the gut microbiome composition. The combination of shared and unique microbial populations across FVB and Wnt1 groups may reflect variations in age and genetic backgrounds.

To identify specific microbial taxa that were significantly impacted by alcohol exposure, we performed LEfSe and two-way ANOVA analysis. These analyses revealed that a number of selective microbial taxa were increased following alcohol exposure; they include the genera *Bacteroides*, *Faecalibaculum* and *Turicibacter,* the species *P. goldsteinii* and *A. muciniphila*. The genus *Bacteroides* is among the most abundant genera in the gut, including species that play diverse roles ranging from beneficial to pathologic (Wexler 2007). For example, *Bacteroides fragilis*, often isolated from extra-intestinal infections, can cause inflammation when it translocates from gut to other organs due to a compromised intestinal barrier (Akhi, Ghotaslou et al. 2015, Akhi, Ghotaslou et al. 2015) (Sun, Zhang et al. 2019). *Bacteroides fragilis* and other *Bacteroides* species such as *Bacteroides uniformis* and *Bacteroides vulgatus* are known to produce beta-glucuronidase, an enzyme involved in the metabolism of estrogens and may influence estrogen-sensitive breast cancer (Fuhrman, Feigelson et al. 2014, Fernández-Murga, Gil-Ortiz et al. 2023). Interestingly, *Bacteroides fragilis* has also been shown to exert anti-cancer and anti-proliferative effects in mouse breast cancer models (Karami, Goli et al. 2023). The genus *Faecalibaculum*, from the *Erysipelotrichaceae* family, plays a critical role in the anti-tumor effects of combined therapies using anti-PD-1 antibody and dietary supplement fucoidan in a breast cancer mouse model (Li, Dong et al. 2024). One of *Faecalibaculum* species, *Faecalibaculum rodentium*, originally identified as anti-tumorigenic in a mouse model for colorectal cancer (Zagato, Pozzi et al. 2020), was shown to counteract the antibiotic-induced tumor growth acceleration in multiple breast cancer mouse models (McKee, Kirkup et al. 2021). Meanwhile, *Turibacter*, typically present at low or moderate levels in gut, is involved in the metabolism of lipids and bile acids (Lynch, Gonzalez et al. 2023). However, higher levels of *Turibacter* have been detected in the intra-tumoral microbiome of patients with triple-negative breast cancer (TNBC) (Wang, Qu et al. 2023) and in the gut microbiota of premenopausal breast cancer patients (He, Liu et al. 2021). *P. goldsteinii* is a low-abundant probiotic in the gut microbiome that supports intestinal integrity and reduces inflammation in conditions such as obesity (Wu, Lin et al. 2019) and pulmonary diseases (Lai, Lin et al. 2022). Finally, *A. muciniphila,* a mucin-degrading bacterium with anti-inflammatory properties, has been associated with improved outcomes in metabolic disorders, intestinal inflammation and several cancers (Cani, Depommier et al. 2022).

The LEfSe and two-way ANOVA analyses also revealed that alcohol exposure led to the reduction of several key microbial taxa: the species *L. intestinalis*, the genera *Candidatus Saccharimonas*, *Herbinix* and *Anaeroplasma*. *L. intestinalis* is a known probiotic that has been shown to support gut health and immune regulation in a variety of disease models (Lim, Song et al. 2021, Lin, Chueh et al. 2023, Wang, Jia et al. 2023). *Candidatus Saccharimonas,* a genus within the family *Candidatus Saccharimonadaceae,* has been identified primarily through genetic analysis but remains uncultured, therefore further research is needed to elucidate its functional role. *Herbinix*, though less studied than other genera in the *Lachnospiraceae* family, is part of a group known for SCFA production, which plays a crucial role in maintaining gut health. This genus is frequently identified in microbiome studies through sequencing data and contributes to the overall microbial composition of the gut (Maus, Bremges et al. 2017, Jia, Yun et al. 2021). *Anaeroplasma* has been shown to exhibit anti-inflammatory effects in lung diseases (Wang, He et al. 2024) and improve bile acid metabolism and enhance gut barrier function in metabolic disorders (Wu, Kaicen et al. 2023, Forlano, Martinez-Gili et al. 2024). Nonetheless, *Anaeroplasma* has been negatively correlated with the efficacy of naringenin, a flavonoid compound in citrus fruits, in the treatment of non-alcoholic fatty liver disease (Cao, Yue et al. 2023).

In summary, our findings demonstrate that alcohol exposure significantly reduces gut microbiome diversity and alters specific microbial taxa. These results underscore the potential role of gut dysbiosis in alcohol-induced tumor promotion and pave the way for further exploration of microbial contributions to mammary tumorigenesis. For instance, taxa such as *Turicibacter*, *Faecalibaculum*, *P. goldsteinii*, *L. intestinalis*, and *A. muciniphila* are known to regulate inflammation, a critical factor in carcinogenesis. Additionally, *Bacteroides fragilis* and other species, including *Bacteroides uniformis* and *Bacteroides vulgatus*, are involved in estrogen metabolism, which may influence the initiation and progression of hormone-related breast cancers. Exploring the involvement of these bacteria in alcohol-driven mammary carcinogenesis represents a promising avenue for future research.

## Data availability statement

The original data presented in the study are available upon request to the corresponding authors. We need to upload data on NCBI server

## Ethics statement

The animal study was approved by the Institutional Animal Care and Use Committee (IACUC) of the University of Iowa. The study was conducted in accordance with the local legislation and institutional requirements.

## Funding

The author(s) declare financial support was received for the research, authorship, and/or publication of this article. This research was supported by NIH grants (AA017226 and AA015407).

## Conflict of interest statement

The authors declare that the research was conducted in the absence of any commercial or financial relationships that could be construed as a potential conflict of interest.

## Notes

### Competing Interest Statement

The authors have declared no competing interest.

