## Supplementary figures and images for "Effects of alcohol on gut microbiome in adolescent and adult MMTV-Wnt1 mice"

### Supplementary Figure 1

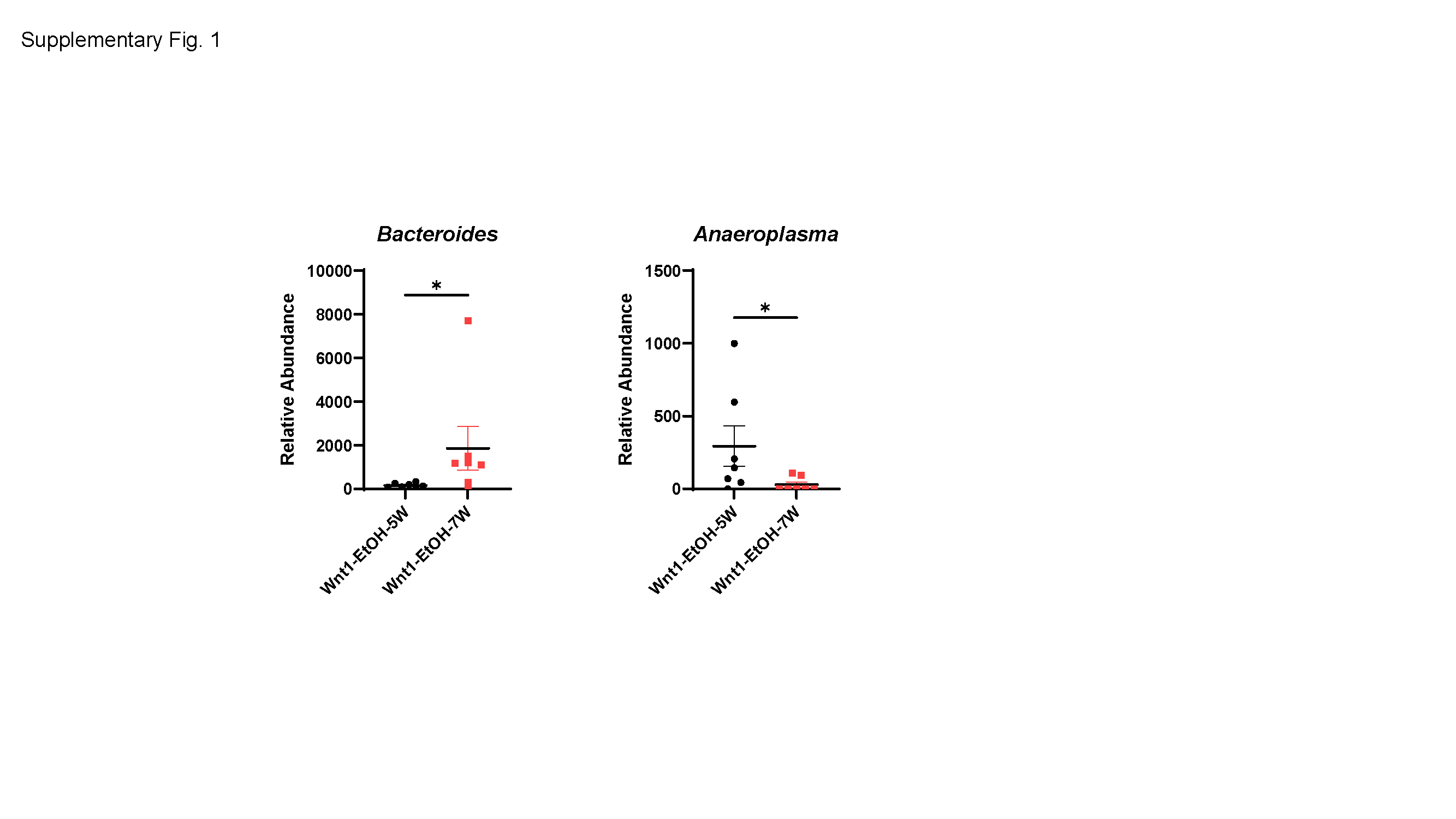

### Supplementary Figure 2

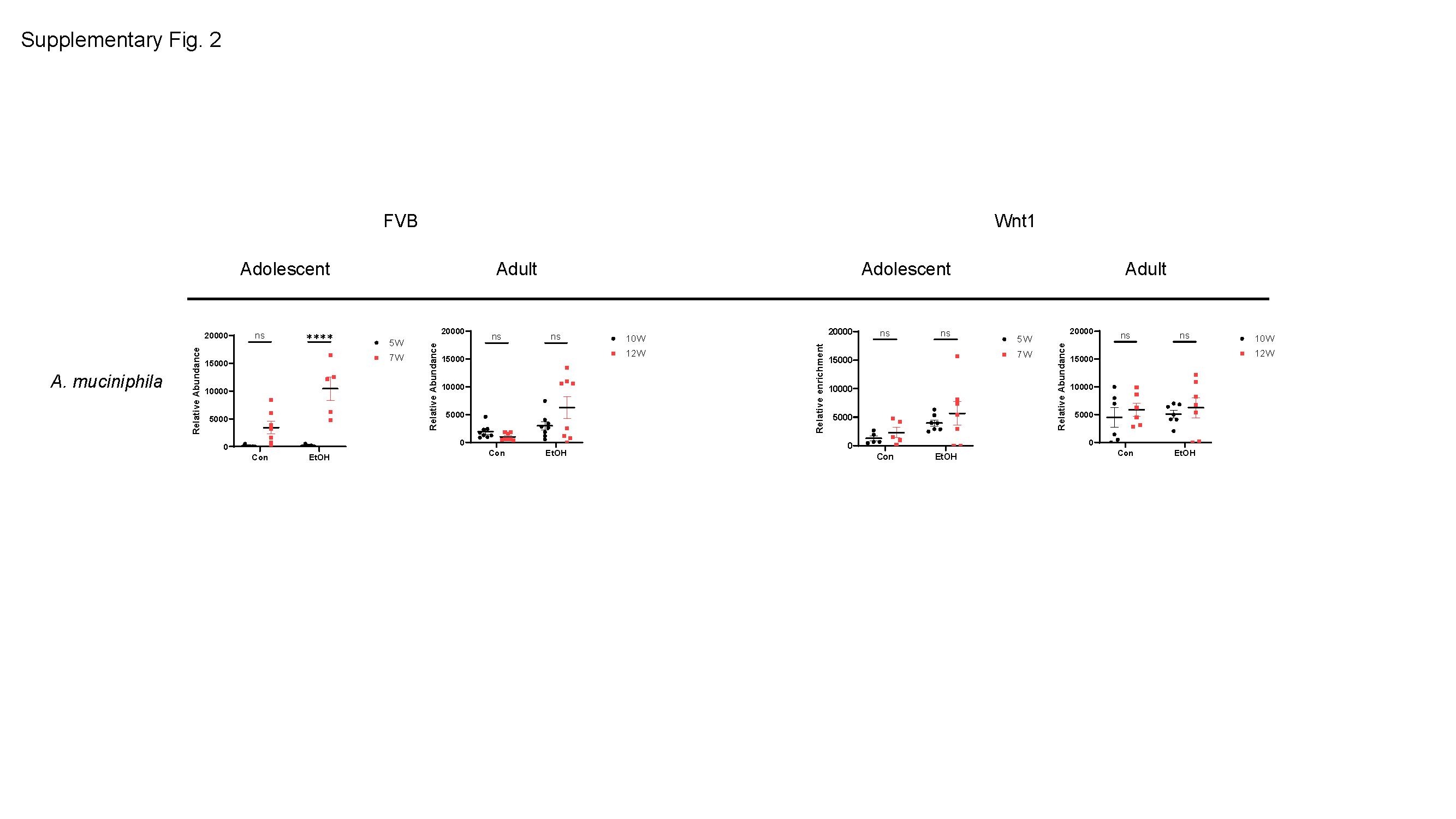
